# Systems pathology analysis identifies neurodegenerative nature of age-related retinal diseases

**DOI:** 10.1101/248088

**Authors:** Tiina Öhman, Fitsum Tamene, Helka Göös, Sirpa Loukovaara, Markku Varjosalo

**Affiliations:** Institute of Biotechnology and Helsinki Institute of Life Science, University of Helsinki, Viikinkaari 1, P.O. Box 65, FI-00014 Helsinki, Finland; University of Helsinki and Unit of Vitreoretinal Surgery, Department of Ophthalmology, University of Helsinki and Helsinki University Hospital, Haartmaninkatu 4 C, FI-00290 Helsinki, Finland

**Author notes:** equal contribution (Sirpa Loukovaara, Markku Varjosalo). **Corresponding authors**: Markku Varjosalo, PhD, Sirpa Loukovaara, MD, PhD.

**Keywords:** epiretinal membrane, macular hole, vitreous humor, MS-based quantitative proteomics, neurodegeneration

## Abstract

Aging is a phenomenon associated with profound medical implications. Idiopathic epiretinal membrane (iEMR) and macular hole (MH) are the major vision-threatening vitreoretinal diseases affecting millions of aging people globally, making these conditions an important public health issue. The iERM is characterized by fibrous tissue developing on the surface of the macula, leading to biomechanical and biochemical macular damage. MH is a small breakage in the macula associated with many ocular conditions. Although several individual factors and pathways are suggested, a systems pathology level understanding of the molecular mechanisms underlying these disorders is lacking. Therefore, we performed mass spectrometry based label-free quantitative proteomics analysis of the vitreous proteomes from patients with iERM (n=26) and MH (n=21) to identify the key proteins as well as the multiple interconnected biochemical pathways contributing to the development of these diseases. We identified a total of 1014 unique proteins, of which many were linked to inflammation and complement cascade, revealing the inflammational processes in retinal diseases. Additionally, we detected a profound difference in proteomes of the iEMR and MH compared to the non-proliferative diabetic retinopathy. A large number of neuronal proteins were present at higher levels in iERM and MH vitreous, including neuronal adhesion molecules, nervous system development proteins and signalling molecules. This points toward the important role of neurodegeneration component in the pathogenesis of age-related vitreoretinal diseases. Despite of marked similarities, several unique vitreous proteins were identified in both iERM and MH conditions, providing a candidate targets for diagnostic and new therapeutic approaches. Identification of previously reported and novel proteins in human vitreous humor from patient with iERM and MH provide renewed understanding of the pathogenesis of age-related vitreoretinal diseases.

## Introduction

Population aging is a global phenomenon with profound medical implications. Tissue dysfunction associated with aging affects all vital organs, including eyes. Various ocular structures are affected by aging, such as the macula, the functional center of the retina responsible for precise central vision. Idiopathic epiretinal membrane (iEMR) and macular hole (MH) are the major vision-threatening vitreoretinal diseases affecting millions of aging people globally, making these conditions an important public health issue^1^. iERM is characterized by the growth of fibrocellular tissue along the inner surface of the retina^2^. Ultimately, impaired repair and excess of fibrosis in iERM eyes leads to biomechanical and biochemical macular damage, development of retinal surface wrinkling with or without shallow tractional retinal detachment, macular vascular distortion, breakdown of the blood–retinal barrier at the retinal pigment epithelial (RPE) level, and vascular leakage. MH, however, is a full-thickness defect of retinal tissue involving the anatomic fovea^3^. Originally, MH was described in the setting of trauma, but it has been associated with many ocular conditions and the great majority of MH cases are idiopathic.

The exact pathogenic mechanisms underlying these conditions are still not known. Older age and the development of anomalous posterior vitreous detachment (PVD) are the generally accepted non-modifiable key risk factors in iERM pathogenesis^4,5,6^. In addition, a number of inflammatory and immunomodulatory processes, chronic oxidative insult, proteolysis, and cytoskeleton remodelling have been implicated in its formation^7,8,9^. Metabolically, retina is the most oxygen consuming tissue in the human body^10^. Therefore, also metabolic stress and altered microvascular retinal blood flow, together with genetic and life-style related factors, such as smoking, could play a role in iERM formation^11,12^. The pathogenesis of idiopathic age-related MH remains unclear, despite a list of theories, and systems level understanding of the disease would be instrumental in developing therapeutic approaches. Currently, pars plana vitrectomy remains the primary treatment option for achieving MH closure and improvement and/ or stabilization of visual acuity in iERM eyes.

The fundamental cell types involved in iERM are retinal pigment epithelium (RPE), Müller cells, astrocytes and microglia that begin proliferating and migrating onto the surface of the retina^13,14^. Especially, microglia, the main retinal immune cells (macrophages), play a key role both in degenerative and inflammatory retinal diseases^15^. Also other cell types present at the vitreoretinal interface, such as hyalocytes, may contribute to the ERM contraction^16^.

The protein composition of vitreous humor is vital for its homeostasis. In healthy eye the homeostasis of retinal extracellular matrix (ECM) is tightly regulated, being altered in ocular disorders, offering a means of indirectly studying the events that take place at the retina^17,18^. Mass spectrometry (MS) based quantitative proteomics provides means for determination of global proteome changes in tissue and cellular level, enabling a molecular level characterization of the complex eye disorders` pathophysiology. Currently, most proteomic studies characterizing disease induced vitreous proteome changes have focused on proliferative and non-proliferative diabetic retinopathy^19^, proliferative vitreoretinopathy^20,21^ and age-related macular degeneration (AMD)^22^, whereas iERM and MH remain less-studied^7,23^.

Therefore, for obtaining in-depth and global understanding on the complex and multi-factorial molecular pathomechanisms underlying these two most typical age-related vitreoretinal interface eye disorders, we performed liquid chromatography - mass spectrometry (LC-MS) based label-free quantitative proteomics analysis of the vitreous proteomes from patients with iERM (n=26), and MH (n=21). The results were compared to the proteomes from diabetic retinopathy patients with macular edema (DME, n=7). We identified a total of 1014 unique proteins, of which the most were linked to inflammation and complement cascade, revealing the inflammation processes in these retinal diseases. We show here that age-related iERM and MH vitreous proteomes differ clearly from DME proteome. A large number of neuronal proteins were present at higher levels in iERM and MH proteomes, including neuronal adhesion molecules, nervous system development proteins and signalling molecules. Bioinformatic analysis revealed that neurodegeneration rather than neuroinflammation seems to play an important role in the pathogenesis of age-related vitreoretinal diseases. Despite of marked similarities, we identified several vitreous proteins that differ between the iERM and MH condition, providing a candidate target for diagnostic and therapeutic approaches. Primary prevention of retinal fibrosis is an important goal. Currently, there is no known way to pharmacologically impact this process, making MS-related investigation an important method to shed more light on the role of various proteins in this harmful process.

## Materials and Methods

### Patients

The study was conducted according to the Declaration of Helsinki and was approved by the Institutional Review Board of Helsinki University Central Hospital, University of Helsinki, Finland. Signed informed consent was obtained from each participant before sampling. Confidentiality of the patient records was maintained when the clinical data were entered into a computer-based standardized data entry for analysis.

Patients were admitted for primary vitrectomy for management of iERM, MH and diabetic retinopathy with macular edema (DME) pathologies in the Unit of Vitreoretinal Surgery, Helsinki University Central Hospital, Helsinki, Finland. Diagnosis and detection of morphologic retinal pathological changes of each studied eye was confirmed using optical coherence tomography (OCT) prior to surgery. OCT (Stratus OCT; Zeiss) or spectral domain ‐ OCT OptoVue RTVue V.5.1 device (OptoVue Inc.) were acquired under mydriatic circumstances. The scan pattern used on Optovue RTVue was a standardized macular protocol, retina MM6, which gives accurate thickness measurements in fovea, perifovea (< 3 mm), and parafovea (< 6 mm). The measurements of central retinal thickness in the innermost foveal area were recorded and analyzed pre- and post-operatively.

Exclusion criteria among iERM and MH patients were vitreous haemorrhage and inflammation, other retinal inflammatory or retinal vascular disorders (retinitis, choroiditis, uveitis, retinal vein occlusion), retinal detachment (RD), AMD or previous ocular trauma. One eye out of the iERM group was excluded because of previous systemic borreliosis as well as one eye out of MH group because of previous glaucoma surgery (trabeculectomy). A total of 54 eyes of 54 patients underwent primary transconjunctival microincision vitrectomy for iERM (n=26), MH (n=21), and DME, n=7 between 2006 and 2016. The clinical systemic and ocular characteristics of the study patients are given in Supplementary Table 1.

### Vitreous sample collection

Undiluted vitreous samples (up to 1000 μl) were collected at the start of the standard 3-port pars plana vitrectomy (25 or 23 Gauge, Constellation Vision System, Alcon Instruments, Inc.) without an infusion of artificial fluid. The samples were collected by manual aspiration into a syringe via the vitrectomy with the cutting function activated. Samples were transferred into sterile microcentrifuge tubes and immediately frozen at -70 °C until laboratory analysis.

In vitrectomy, if the vitreous was attached to the posterior retina, posterior vitreous detachment was induced by suction with the vitrectomy probe over the optic disc. To visualize and identify the posterior hyaloid, epiretinal membranes and the internal limiting membrane, intravitreal vital dyes were used (chromovitrectomy). Diluted indocyanine green, MembraneBlue-Dual^®^ (D.O.R.C., Zuidland, The Netherlands) or ILM-blue^®^ (D.O.R.C. Zuidland, The Netherlands) dye-assisted ERM ± ILM or plain ILM peeling was performed. Peeling of ERM and ILM was carried out using a pinch-peel technique with fine-tipped forceps. In all MH eyes a fluid-air exchange was performed with subsequent gas exchange.

### Sample preparation for the LC-MS

The collected patient vitreous samples were centrifuged at 18 000 x g for 15 minutes at 4°C to clear the samples from cell debris. Average protein concentrations (mg/ml) of the vitreous samples were determined using bicinchoninic acid (BCA) protein assay kit (Pierce, Thermo Scientific) according to the manufacturer’s instructions. 100 μg of total protein per sample was taken for LC-MS analysis. UREA was added to the individual samples (to final concentration 1M), the proteins reduced with tris(2-carboxyethyl)phosphine (TCEP) and alkylated with iodoacetamide. The proteins were then digested to peptides with Sequencing Grade Modified Trypsin (Promega). The resulting tryptic peptides were purified with C18 microspin columns (Nest Group, Southborough, MA, USA) before subjected to LC-MS/MS analysis.

### LC-MS/MS Analysis

The LC-MS/MS analysis was done with a Q Exactive ESI-quadrupole-orbitrap mass spectrometer coupled to an EASY-nLC 1000 nanoflow LC (Thermo Fisher Scientific), using the Xcalibur version 3.1.66.10 (Thermo Scientific) essentially as described (Loukovaara et al. 2015). The tryptic peptide sample mixture was automatically loaded from autosampler into a C18-packed precolumn (Acclaim PepMap™100 100 μm × 2 cm, 3 μm, 100 Å, Thermo Scientific) in 10 μl volume of buffer A (1 % acetonitrile, 0.1 %, formic acid). Peptides were transferred onward to C18-packed analytical column (Acclaim PepMap™100 75 μm × 15 cm, 2 μm, 100 Å, Thermo Scientific) and separated with 120-minute linear gradient from 5 to 35 % of buffer B (98 % acetonitrile, 0.1 % formic acid) at the flow rate of 300 nl/minute. This was followed by 5 minute gradient from 35 to 80 % of buffer B, 1 minute gradient from 80 to 100 % of B and 9 minute column wash with 100 % B at the constant flow rate of 300 nl/minute. The mass spectrometry analysis was performed in data-dependent acquisition in positive ion mode. MS spectra were acquired from *m/z* 200 to *m/z* 2000 with a resolution of 70,000 with Full AGC target value of 1,000,000 ions, and a maximal injection time of 100 ms, in profile mode. The 10 most abundant ions of which charge states were 2+ to 7+ were selected for subsequent fragmentation (higher energy collisional dissociation, HCD) and MS/MS spectra were acquired with a resolution of 17,500 with AGC target value of 5000, a maximal injection time of 100 ms, and the lowest mass fixed at *m/z* 120, in centroid mode. Dynamic exclusion duration was 30 s.

### MS1 Quantification and Protein Identification

The Progenesis LC-MS software (v4.1, Nonlinear Dynamics Limited, Tyne, UK) was used to obtain the MS1 intensities of peptides for label-free quantification. The run with the greatest similarity to all other runs was automatically selected as the alignment reference. All runs were then aligned to a reference run automatically and further adjusted manually. Retention time was limited to 10-130 minutes excluding first and last ten minutes of the recorded data. Only peptides with charge states from +2 to +7 were allowed. For protein identification the MS/MS-scan data acquired from Progenesis LC-MS was searched against the human component of the UniProtKB- database (release 2016_06 with 20154 entries) using SEQUEST search engine in Proteome Discoverer™ software (version 1.4, Thermo Scientific). Carbamidomethylation (+57.021464 Da) of cysteine residues was used as static modification and oxidation (+15.994491 Da) of methionine was used as dynamic modification. Precursor mass tolerance and fragment mass tolerance were set to less than 15 ppm and 0.05 Da, respectively. Maximum of two missed cleavages were allowed. Results were filtered to a maximum false discovery rate of 0.05. For protein identification, peptide spectrum match was ≥ 2.

To validate the sample reproducibility, on feature alignment and detection level, five samples were analysed in technical replicates. The distribution of the MS1 feature alignment and MS1 quantitation level correlation showed extremely high reproducibility (Supplementary Table S2).

### Dot Blot Analysis

5 μg of total protein from the vitreous samples were dot blotted to nitrocellulose membrane using Bio-Rad 96-well dot blot system (Bio-Dot^®^ Microfiltration Apparatus, Bio-Rad) according manufactures instructions. Six antibodies against the differentially expressed proteins between samples were selected based on availability of high quality antibodies suitable for Western Blotting. Signals were visualized using Amersham ECL Western Blotting analysis system (GE Healthcare, UK). The dot blots were analysed and quantified using Dot Blot Analyzer for ImageJ30. For each of the proteins, the Pearson´s correlation was calculated between the corresponding MS1 and dot blot intensities.

### Data Processing

Statistical significant test of abundance changes between iERM, MH and DME eyes were conducted using Student’s t-test. An abundance change with q-value of 0.05 or less was considered as significant change. Q-values were used instead of conventional p-values to maximize the power of statistical test. The hierarchical clustering of the identified proteins was constructed using Gene Cluster 3.0. The line graph and boxplots were produced using R version 3.3.2. Gene Ontology (GO) annotations were obtained from DAVID bioinformatics resources^24,25^. The cellular locations of identified proteins, being either intracellular, transmembrane or extracellular, were extracted from Phobius predictor^26^. The MediSapiens database (www.medisapiens.com) was used to study the gene expression levels across healthy human tissues^27^. For obtaining the known protein-protein interactions the PINA2 protein interaction database was used^28^.

### Data Availability

The peptide raw data have been uploaded to the MassIVE public repository (http://massive.ucsd.edu.), the MassIVE ID MSV000081839.

## Results

### Study plan and the patients’ pre-operative analyses

iERM and MH are complex age-related vitreoretinal interface diseases. Although both are related to aging and have similar visual disturbances and symptoms including metamorphopsia, photopsia, blurred vision, and decreased visual acuity, iERM and MH are different pathological conditions. iERM displays as thin layer of scar tissue that has formed on the posterior pole, at human anatomic macula, whereas MH manifests as a partial or full thickness loss of tissue in the central retina (Fig. 1A). An optical coherence tomography (OCT) scan through the fovea of the iERM eye reveals the abnormal organization of the retinal layers including epiretinal fibrosis and secondary cystic macular edema (Fig. 1B).

**Fig. 1.**
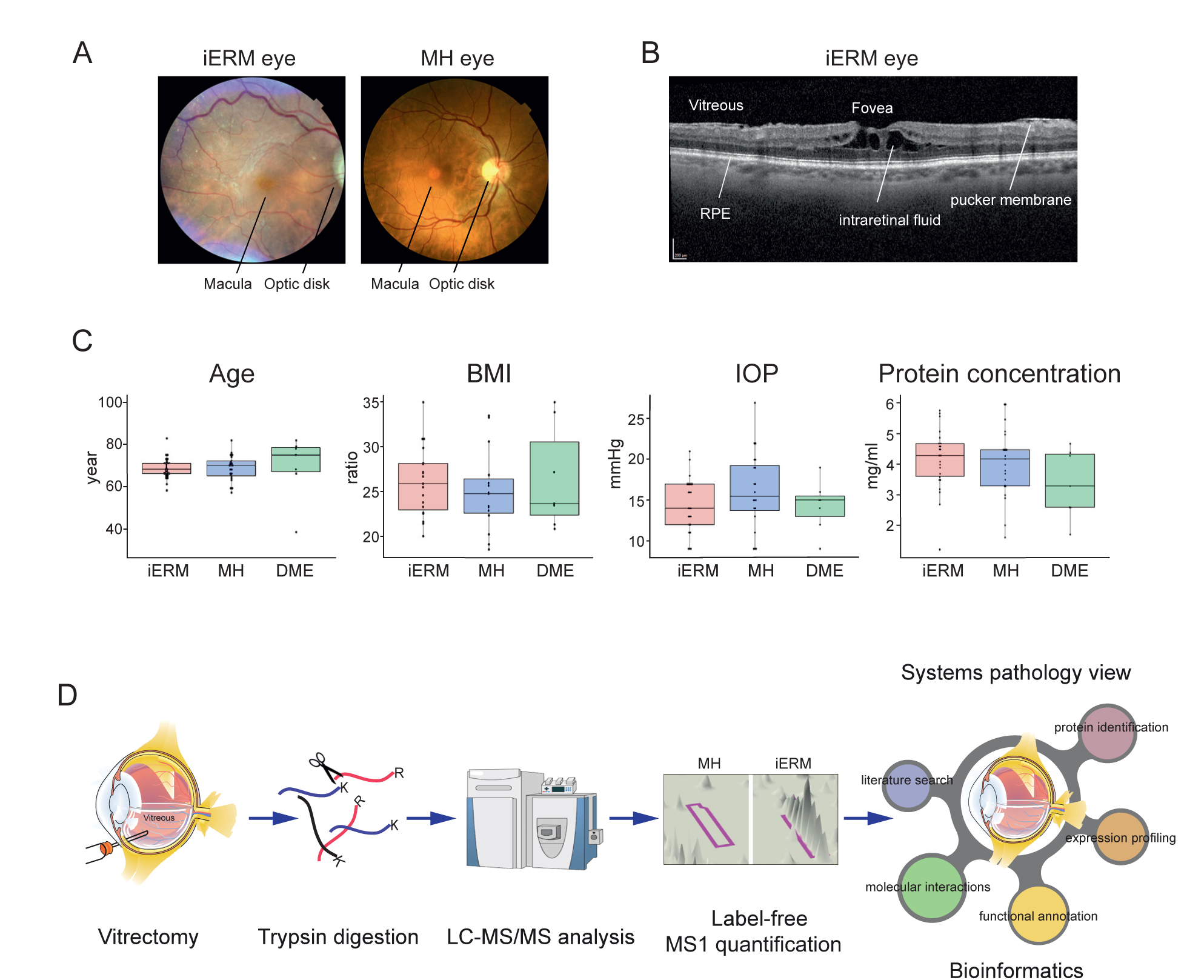
Patient characterization and quantitative proteomics pipeline. A) Fundus photograph of the central and peripheral retina, optic disk and macula from patients with iERM or MH. The macula is located in the posterior pole of the eye. In the center of the macula, a shallow depression in the retina (the fovea) marks the area with the highest visual acuity. B) An optical coherence tomography (OCT) scan through the fovea of the iERM eye reveals the abnormal organization of the retinal layers including epiretinal fibrosis and secondary cystic macular edema. Key: RPE = retinal pigment epithelial cells, scale bar 200 μm. C) The demographics of the iERM, MH and DME patients, showing the distribution of the age, body mass index (BMI), preoperative intraocular pressure (IOP) and protein concentration (mg/ml). D) The experimental workflow used for identified and quantified human vitreous proteins from patients with iERM, MH or DME. Vitreous samples were collected in vitrectomy, proteins extracted and digested with trypsin and the resulting peptides were analysed via LC-MS/MS. The label-free quantification was done using Progenesis LC-MS analysis software and the protein identification using SEQUEST search engine. Bioinformatics approaches were used to combine our proteome data with the existing knowledge in order to obtain a systems pathology view on the differences of the molecular ethiologies of these eye diseases.

The transparent collagenous human vitreous is in close contact with the retina and lacks its own vasculature. Because of this close interaction, the physiological and pathological conditions of the retina are reflected directly on the protein composition of the vitreous. In order to get an in depth view of the vitreous humor proteome in the aging human eye, we collected and analysed a total of 54 vitreous humor patient samples; 26 were obtained from iERM and 21 from MH patients. In addition, seven age-matched type 2 diabetic retinopathy patients with macular edema (DME) were included in analysis as a control group. Patients with prior surgical complications (such as cataract complications) were not enrolled to this study. The mean age of iERM, MH and DME patient groups was highly similar 68.7 ± 5.0, 68.6 ± 6.2 and 69.4 ± 15.0 years, respectively (Fig. 1C). Neither body mass index (BMI) proportion nor intraocular pressure (IOP) varied significantly between the three sample groups (Fig. 1C). The average protein concentration in the vitreous samples of the iERM, MH and control-DME were 4.2 ± 1.0, 4.1 ± 1.2 and 3.4 ± 1.1 mg/ml, respectively (Fig. 1C). Detailed patient demographics are shown in Supplementary Table S1.

### The analysis of the patients´ vitreous humor proteomes

Vitreous samples from each patient were cleared from possible insoluble cellular fractions and analysed for proteome composition. The proteins, digested into peptides, were analysed with LC-MS/ MS and the corresponding protein identity and abundance were obtained with label-free quantification of the MS1 spectra with Progenesis LC-MS software in combination with SEQUEST search engine (Fig. 1D). One sample from MH group was chosen as a reference run, to which all the other runs were aligned. The median alignment percentage for MH samples was 80.7%, showing high similarities between the samples (Fig. 2A). The median alignment percentages for iERM and DME sample groups compared to MH reference were 67.6% and 52.1%, respectively, indicating differences in vitreous proteomes of iERM and DME patients compared to the MH patients.

**Fig. 2.**
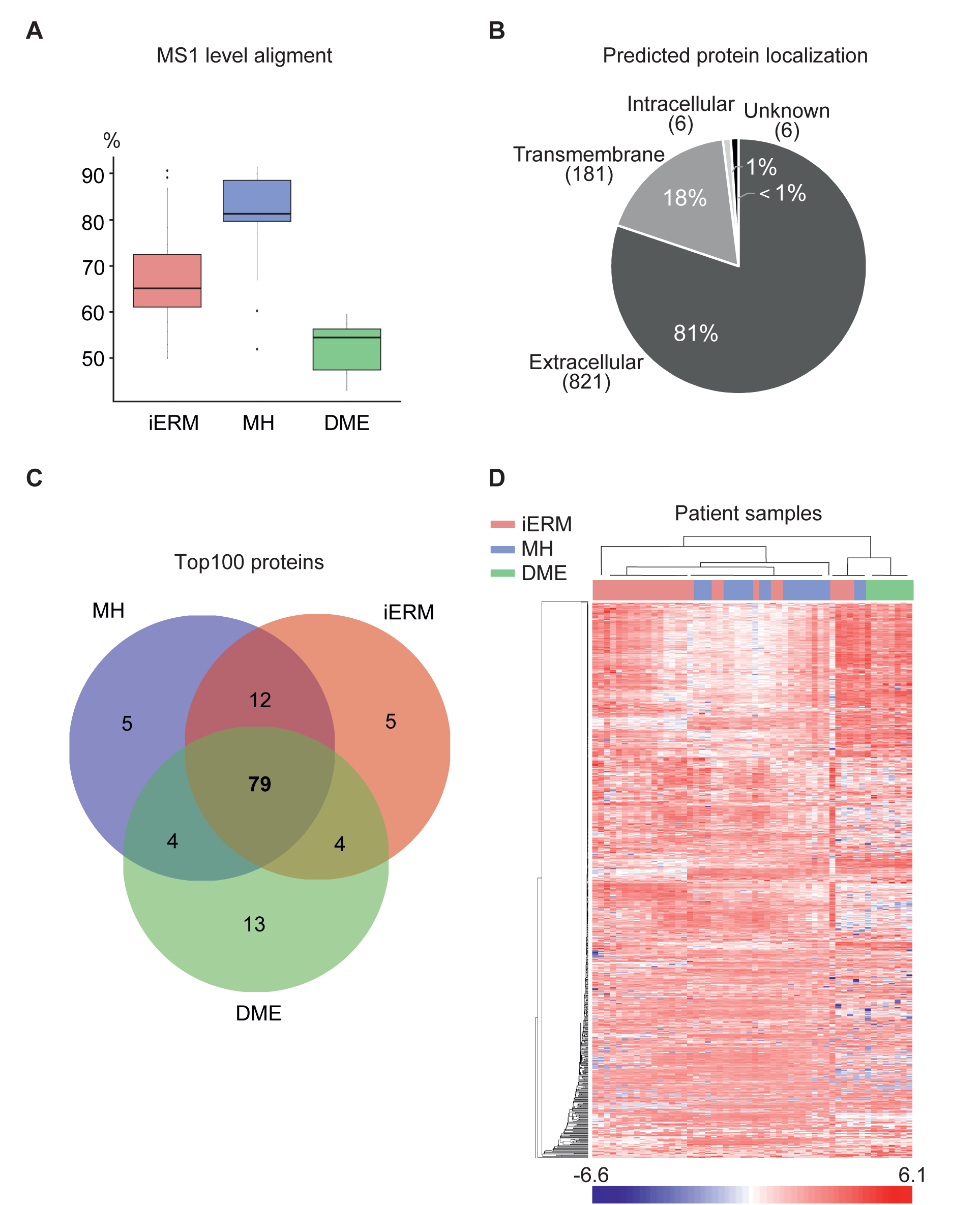
The vitreous proteomes from patients with iERM, MH or DME show differential composition between the diseases. A) The MS1 spectra alignment percentages for iERM, MH and DME sample MS analysis runs compared to MH reference run. B) Cellular localizations of the detected proteins were predicted using Phobius predictor software and found predominantly to be either extracellular or transmembrane. C) Venn diagram of 100 the most abundant protein in iERM, MH and DME sample groups shows good overlap between the disease groups, whereas D) the disease groups separate well after hierarchical clustering of the global quantitative proteomes. Red in the heatmap indicates high while blue denotes low log2-normalized MS1 intensities.

On the protein level, we could identify in total 1014 unique proteins with 4323 nonconflicting peptides (Supplementary Table S3). To assess the cellular localization of the identified proteins (Fig. 2B, Supplementary Table S3) we used Phobius software^26^, and more than 99% of intravitreal proteins were identified either as extracellular or having transmembrane domain. This corresponds well with the fact that most vitreous proteins are expected to be secreted or shed from the surrounding tissues^8,19^.

Of the 1014 identified proteins, 934 were quantified with average MS1 intensity over 1×10^4^ at least in one of the sample groups (Supplementary Table S4; MH, iERM or DME). The most common proteins in each sample group were highly similar (Fig. 2C), consisting of high abundance proteins such as serum albumin, transferrin, complement factors and apolipoproteins. To profile the vitreous proteomes, or more precisely their possible quantitative differences, we performed hierarchical clustering analysis of the 934 quantified proteins (Fig. 2D). In spite of high inter-group similarity on the level of the most abundant proteins, the clustering revealed three large separated protein clusters, defining the differential protein changes induced by iERM, MH and DME diseases.

To further validate MS data, we performed a high-throughput dot blot analysis for 6 proteins; the dot blot results correlate highly with MS1 data (median 0.65, Supplementary Fig. 1). The high correlation between dot blot analysis and MS1 quantification of vitreous samples has also been shown previously^19^.

### DME vitreous proteomes differ from iERM and MH proteomes

When the iERM and MH proteomes were compared to the vitreous proteome of DME, a clear difference was detected among the groups (Fig. 3A). In the iERM proteome 80 proteins were present at higher level and 131 proteins at lower level compared to DME proteome (the abundance ratio >2, q-value < 0.05, Supplementary Table S5). Likewise, in the MH proteome 174 proteins were present at higher level and 123 proteins at lower level compared to DME proteome (Fig. 3A, Supplementary Table S6). When the differently expressed proteins were classified according to their involvement in different biological processes using DAVID bioinformatic resource^24,25^, we observed that the more abundant proteins in the DME group belonged to a limited number of biological processes, mainly immune system. More specifically, the proteins were associated with blood coagulation, fibrinolysis and platelet aggregation linked to wound healing, as well as complement activation and phagocytosis linked to inflammatory processes (Fig. 3B). This observation is consistent with our previous report showing that inflammation and complement cascade play a significant role in diabetic retinopathy and especially in the most advanced proliferative form^19^. Many proteins involved in the classical and alternative pathways of complement activation were also detected in MH and iERM proteomes, indicating that inflammation may be a common nominator both in fibroproliferative and non-fibroproliferative retinal disease.

**Fig. 3.**
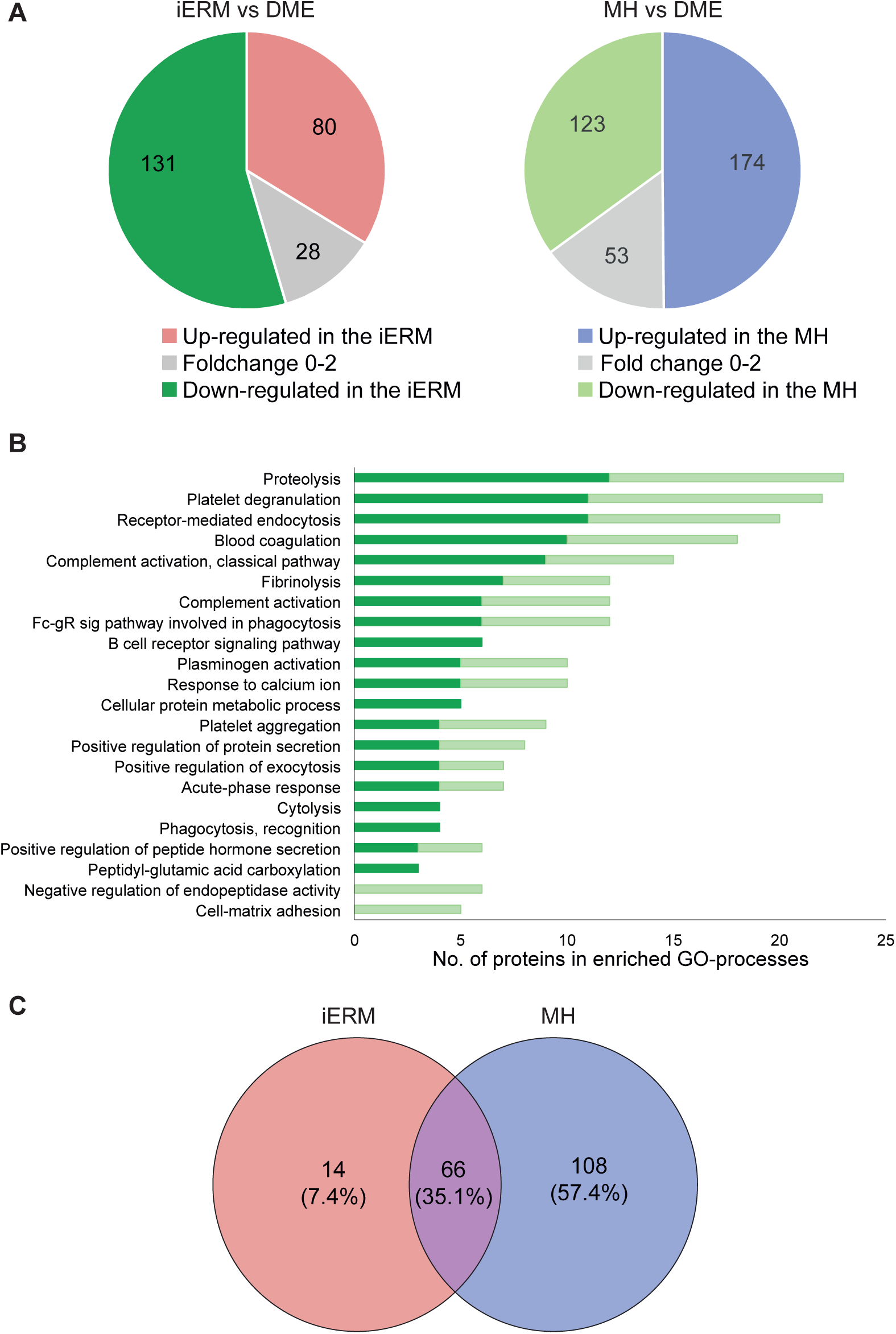
iERM and MH vitreous proteomes differ clearly from DME. A) The abundance of 240 or 351 proteins differed statistically significantly (q-value ≤ 0.05) between iERM and DME (left panel) or between MH and DME groups (right panel), respectively. 80 proteins were present at higher level and 131 proteins at lower level in the iERM proteome (abundance ratio difference >2 fold) and 174 proteins were present at higher level and 123 proteins at lower level in the MH proteome when compared to DME proteome. B) 131 and 123 proteins that were more abundant in the DME group were categorized according to their involvement in biological processes (Gene Ontology, Biological Processes terms) via DAVID bioinformatics resources. Dark green indicates proteins that were upregulated in DME compared to iERM (131) and light green upregulated in DME compared to MH (123). C) Venn diagram of 80 and 174 proteins upregulated in iERM and MH, respectively, shows a marked overlap.

The proteins being upregulated in iERM (n=80) and MH (n=174) showed a marked overlap (66 proteins) (Fig. 3C). Similarly, these two retinal disease conditions shared the most frequently enriched biological processes (Fig. 4A). The common proteins were associated with cell adhesion, movement of cell, nervous system development, cell signaling and proteolysis (Table 1). This finding of multiple proteins being linked to cell adhesion, migration and extracellular matrix supports the hypothesis that the development of macular damage requires cell migration from within the retina and extracellular matrix containing fibrous element formation. Of note, we detected six members of neuronal cadherin and catenin family (CAD12, CADH2, CSTN1, CSTN3, CTNB1, CTND2) that were significantly upregulated in iERM and MH samples (Fig. 4B). Cadherin-catenin complex is the main component of the intercellular adherens junction that contributes both to tissue stability and dynamic cell movements^29^. Cadherin-based adherens junctions are involved in various processes of neuronal development, affecting neuronal progenitor cells including retinal stem cells^30^. In retina, in normal physiological circumstances, cadherin complex contributes to RPE cell stability, whereas in some pathologic circumstances, it facilitates RPE cell motility and migration^31^. In addition, cadherin-catenin complex has a critical role in regulating Wnt signaling^32^, and interestingly, the Wnt signaling pathway was one of the main enriched processes detected in iERM proteomes (Fig. 4A).

**Table 1.**
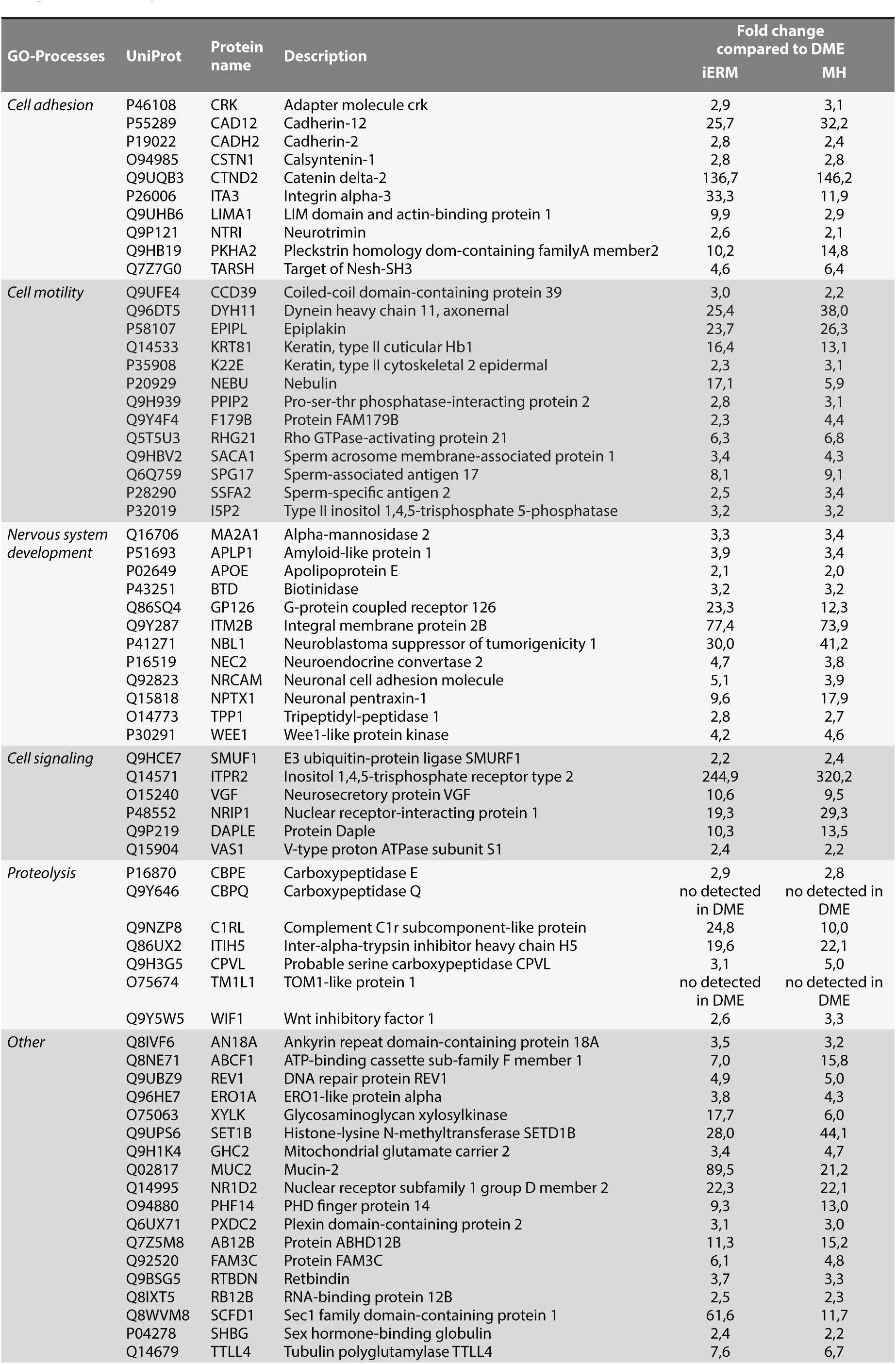
66 common proteins that present in higher level in iERM and MH proteomes, respectively, when compared to DME proteome.

**Fig. 4.**
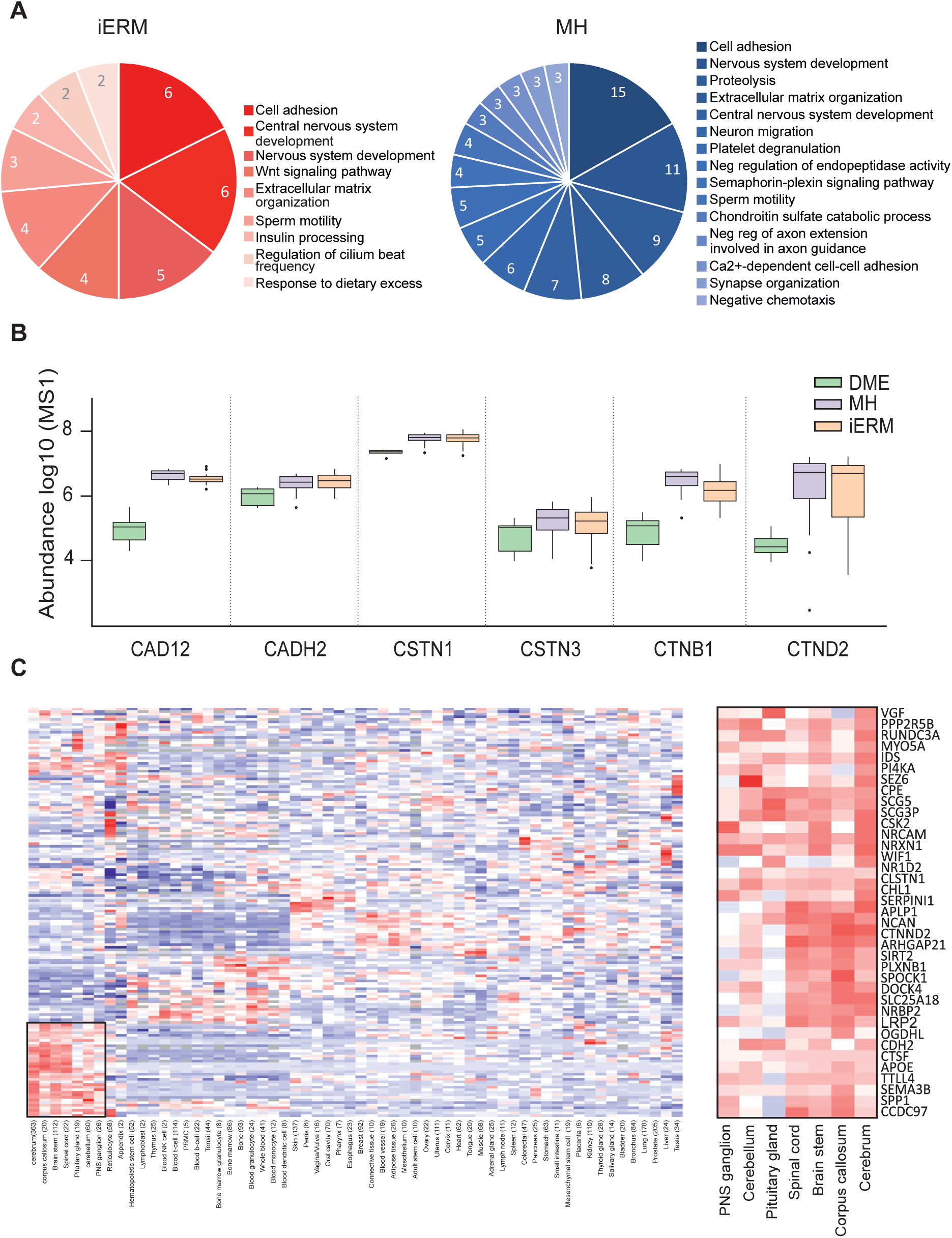
Classification of the proteins up-regulated in iERM and MH sample groups compared to DME group. A) Biological processes associated with the 80 and 174 proteins upregulated in iERM and MH groups, compared to DME group, respectively. B) MS1 intensity (log-scale) of enriched group of adhesion molecules, involved in cadherin-catenin complex, illustrate their abundant presence in iERM and MH C) Hierarchical clustering of the iERM and MH sample upregulated proteins based on their gene expression profiles in healthy human tissues. The iERM and MH upregulated proteins are expressed clearly in separate cluster consisting of, in majority, neuronal tissues.

The other interesting protein groups up-regulated in iERM and MH proteomes were involved in cell motility, including proteins CCD39, DYH11, SPG17 and EPIPL, which regulate cilium movement and wound healing. In retina, photoreceptors have unique sensory cilia that are essential for eye health^33^. The patient with defects in ciliary motility develop retinal degeneration^34^. It has also been shown that primary cilia coordinate a series of signaling pathways and regulate cell migration during development and in wound healing^35^.

Importantly, also specific processes, such as Wnt signaling for iERM samples and semaphorin-plexin pathway for MH samples, were detected in iERM and MH proteomes (Fig 4A). Aberrant Wnt/beta catenin signaling has been implicated in major inflammatory and neurodegenerative disorders^36^ and with a variety of human hereditary diseases. Currently, modulation of Wnt signaling is actively studied in the field of cancer, regenerative medicine and wound healing^37^. However, our understanding of the Wnt pathway is incomplete and many questions in this field remain unanswered.

Semaphorin-plexin pathway is also known to have function in neural and immune systems, as well as in angiogenesis^38^. Intravitreal semaphorin 3A concentration has previously been shown to be increased in patients suffering from diabetic retinopathy^39^. However, in DME proteome in our study, semaphorins, including 3A, were detected only at very low level, whereas several semaphorins (3A, 3B, 4B, 3F, 7A) and their binging partner plexin B1 were present at higher levels in MH proteome, suggesting an important role of semaphorin-plexin signaling in MH, not in DME. Altogether, the role of Wnt or semaphorin-plexin signaling pathway in MH or iERM processes has not been explored in great detail and requires therefore further studies.

### The iERM and MH vitreous proteomes display surprisingly high abundance of neuronal proteins

To assess the cellular origin of the upregulated proteins in iERM and MH, we analyzed the gene expression profiles using the MediSapiens database, which includes the gene expression data from over 3000 samples from 60 “healthy” human tissues (<u>www.medisapiens.com</u>). Only one clear cluster could be detected, showing the accumulation of neuronal origin proteins in the vitreous proteomes (Fig. 4C). To further analyze these neuronal proteins, we used PINA2 database to derive the protein-protein interactions for these proteins (Fig. 5A). According to PINA2 analysis, the 37 up-regulated proteins found in neuronal cluster have altogether 90 interactors that were identified in our vitreous proteome analysis (Fig. 5A, Supplementary Table S7). The interacting proteins formed several protein groups, of which the largest were apolipoproteins, keratins and signaling proteins. Also several proteins involved in neuronal system development and cell adhesion were detected.

**Fig. 5.**
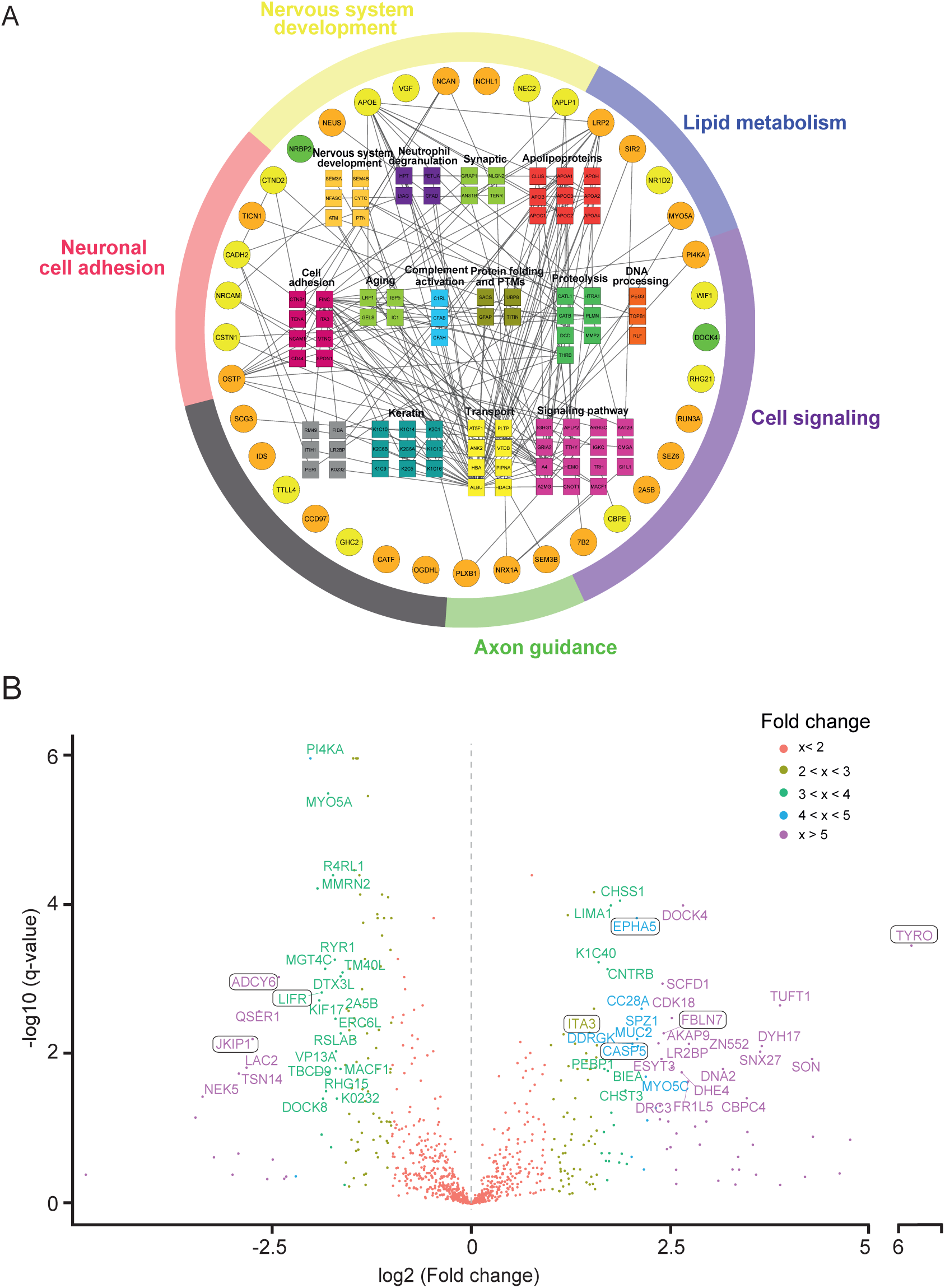
Systems level analysis highlights the interconnectivity of the iERM and MH upregulated proteins and suggests novel biomarker candidates for these diseases. A) Interaction analysis of the differentially abundant neuronal proteins. 37 neuronal proteins that were present at higher level in the iERM and MH proteome were analysed using public PINA2 protein interaction database (green nodes indicate proteins that were upregulated in iERM samples, orange nodes upregulated in MH, and yellow nodes in both samples). Interaction analysis reveals all together 90 interacting proteins found in our vitreous analysis, classified based on their biological processes. B) Comparison between iERM and MH reveals several potential iERM biomarker. Volcano blot analysis of differentially expressed proteins in iERM and MH samples. Short name of proteins were given to proteins with q-value < 0.05 and fold change > 3 (exception ITA3 with fold change 2.8). Potential iERM biomarker candidates are highlighted.

Network analysis showed a central role of apolipoproteins in vitreous proteome (Fig. 5A). Previously, APOA1, APOB, APOH and APOC have been reported as biomarkers for diabetic retinopathy^19,40,41^. However, according to our study, APOE and APOB were more abundant in iERM and MH samples than in diabetic control samples, indicating the specific role of these apolipoproteins in iERM and MH conditions. Therefore, our findings suggest that APOB is not a suitable biomarker for diabetic retinopathy.

Moreover, various keratins have been identified as binding-partners of upregulated neuronal proteins. Generally, keratins are broadly expressed in epithelial cells, but they have also been detected in normal nervous tissue^42^. The detection of multiple different keratins in proteome analysis indicates a specific role of keratins in the age-related eye diseases, even though some keratins can also be contaminants linked to sample preparation. Keratins are a class of intermediate filament proteins that participate in scar formation and wound healing^43^. Since the iERM is a condition where a thin sheet of scar tissue grows on the surface of the retina, keratins could also play a role in iERM process.

Interestingly, many proteins directly associated with neurodegeneration were also detected with higher abundance in the iERM and MH samples than in DME samples, including amyloid-like protein 1 and 2 (APLP1, APLP2), integral membrane protein 2B (ITM2B), amyloid beta A4 protein (A4) and tripeptidyl-peptidase 1 (TPP1). The presence of a member of amyloid precursor protein gene family in vitreous iERM and MH humor suggests that there could be a link between age-related eye diseases and neurodegenerative diseases in the brain. Similar morphological and functional changes in microglial and neuronal activities, such as reported in the brain of Alzheimer´s patient, may also occur in retina^44^. Actually, neurodegenerative processes that have been characterized in central nervous system disorders have also been detected in ocular pathologies, such as glaucoma and age related macular degeneration^45^.

### Comparison of iERM and MH proteomes

The iERM and MH proteomes were highly similar. However, as they clearly are different pathological conditions, we compared the iERM proteome to MH proteome to assess for potential biomarkers of these diseases. Altogether, 160 proteins differed significantly between iERM and MH groups (q-value < 0.05), with 53 proteins being up-regulated in iERM (abundance ratio >2) and 65 proteins in MH samples (Fig. 5B, Supplementary Table S8). Since the differentially expressed proteins could not show any enrichment pathways, these up-regulated proteins iERM and MH groups represent different biological processes, allowing us to select several highly expressed individual proteins that displayed a significantly different abundance in iERM or MH samples, as potential biomarker candidates for diagnosis and prognostic approach.

Tyrosinase (TYRO) is the rate-limiting enzyme oxidase responsible for melanin biosynthesis in the retinal pigment epithelium (RPE) of the eye. RPE cells contain different types of melanin granules^46^ (Boulton, 2014), making them the probable origin of increased tyrosinase. Melanin has an important role in retinal development and protection against light-induced oxidative stress, and melanin levels have been associated with age-related macular degeneration^47,48^. Our analysis showed that tyrosinase levels were 70-fold higher in iERM eyes than in MH eyes and 40-fold higher in iERM eyes than DME eyes, indicating a specific role of tyrosinase in iERM formation.

Fibulin-7 (FBLN7) is a cell adhesion molecule expressed also in the retina. Recently, it was shown to play a role in preventing AMD^49^ (Sardell et al, 2016). In addition, it has been found to negatively regulate angiogenesis^50^, which could be a mechanism for AMD regulation. FBLN7 interacts with beta1 integrin^51^, which forms a heterodimer with alpha3 integrin (ITA3) that was found to be up-regulated protein in our iERM samples. In the eye, integrin receptors have been closely associated with ocular surface inflammation, vitreolysis and choroidal and pre-retinal angiogenesis. α3β1 integrin mediates the attachment of the vitreous to the retinal surface^52^, which make these adhesion molecules a potential therapeutic target for iERM.

Caspases are a family of protease enzymes playing essential roles in programmed cell death and inflammation. During eye development, cell death allows selection of appropriate synaptic connections^53^. Dysregulation of programmed cell death, however, has been linked to the pathogenesis of several retinal diseases, including retinal detachment and AMD^54,55^. Caspase-5 (CASP-5), found to be highly up-regulated in our iERM sample group, is a poorly characterized member of the caspase subfamily. It interplays with caspase-1 in inflammatory responses in RPE cells but the functional roles of CASP-5 in ocular inflammatory diseases are essentially unknown^56^. EPHA5 receptor tyrosine kinase plays also a key role in the development of the eye and visual system^57^, however, the association between EPHA5 and retinal dysfunctions has not been reported.

As regards the MH eyes, three interesting signaling molecules, Janus kinase and microtubule-interaction protein 1 (JAKMIP1), leukemia inhibitory factor receptor (LIFR) and adenylate cyclase type 6 (ADCY6) were presented at significantly higher level in MH proteome than in iERM or DME proteomes, and may therefore be considered as a potential MH marker. JAKMIP1 interacts with Tyk2, a member of Janus kinase family, which has been shown to control survival and proliferation of retinal cells^58^. LIFR, together with its ligand leukemia inhibitory factor (LIF), participates also in neuroprotection by activating an endogenous rescue pathway that protects viable photoreceptor cells^59^. ADCY6 is a member of adenylate cyclase family that synthesizes cyclic adenosine monophosphate or cyclic AMP from adenosine triphosphate. Two members of the family, ADCY1 and ADCY8, work together to facilitate midline crossing of retinal axons, but the role of ADCY6 in retinal function, has not been reported^60^.

## Concluding remarks

Systemic data of intravitreal biochemical factors and signaling pathways related to the age-related retinal diseases iERM and MH are rather scarce. This study utilized label-free quantitative mass spectrometry to investigate the vitreous proteomes in iERM and MH eyes. Precise bioinformatics analysis was used struggling to reveal novel candidate protein groups and signalling pathways involved in iERM and MH formation, potentially guiding the development of further pharmacological treatments or therapies. As a conclusion, our results illustrate the following: (1) iERM and MH are complicated pathological processes, involving inflammation, extracellular matrix dysfunction and fibrosis; (2) the surprisingly large amount of neuronal proteins were significantly more abundant in vitreous proteome from age-related eye diseases iERM and MH than in DME, indicating the neurodegenerative background of these two age-related pathologies; (3) Wnt signalling involved in iERM and semaphorin-plexin pathway in MH formation were novel findings; and (4) the identity of proteins differing significantly between the iERM and MH condition, providing a candidate list of possible targets for diagnostic, prognostic and/or therapeutic approaches

Human vitreous proteomics has rapidly expanded the list of potential protein biomarkers and molecular disease pathways during last decade. Although the differences between iERM pathogenesis compared with MH were actively investigated to provide new potential biomarkers to use for diagnosis and prognostic approach, we could observe several highly upregulated proteins in iERM and MH proteomes that provide further insight into the pathology of these two age-related eye diseases. The vitreous proteome atlas of iERM and MH, generated in this study, will provide plethora of information and from the basis of future diagnostic and therepautic approaches. However, although we were able to provide insight into the mechanisms of these diseases as well as possibly guide the further development of efficient treatments, further studies are required for finding a cure for these pathologies.

## Acknowledgements

Acknowledgements: Publication of this article was supported by grants from the Academy of Finland (nos. 288475 and 294173, M.V.), Sigrid Jusélius Foundation (M.V.), University of Helsinki Three-year Research Grant (M.V.), Biocentrum Helsinki (M.V.), Biocentrum Finland (M.V.), Instrumentarium Research Foundation (M.V.), the Finnish Eye Foundation (S.L.), the Mary and Georg C. Ehrnrooth Foundation (S.L.), and HUCH Clinical Research Grant (TYH2016230, S.L.). We thank Dr. Petri Aaltonen and Dr. Virpi Raivio for collecting individual vitreous samples, and Dr. Petri Auvinen and Dr. Adrian Smedowski for critical reading and comments on the manuscript. Xiaonan Liu is acknowledged for helping with figure 5. The fundus photographs are courtesy of Dr. Petri Tommila.

## Supplementry information

**Supplementary Figure S1:** The correlation between dot blot analysis and MS1 quantification of vitreous samples.

**Supplementary Table S1:** Patient demographics and operated eye characteristics.

**Supplementary Table S2:** Reproducibility of the used analysis pipeline.

**Supplementary Table S3:** The identified vitreous protein with a peptide-spectrum match score (PSM) ≥ 2. The cellular locations of identified proteins, being either intracellular, transmembrane or extracellular, were extracted from Phobius predictor.

**Supplementary Table S4:** Normalized MS1 intensities of the 934 quantified proteins.

**Supplementary Table S5:** Significantly differed proteins between iERM and DME proteomes, q-value < 0.05. 80 proteins were present at higher level (red) and 131 proteins at lower level (yellow) in the iERM proteome when compared to DME proteome, the abundance ratio >2.

**Supplementary Table S6:** Significantly differed proteins between MH and DME proteomes, q-value < 0.05. 174 proteins were present at higher level (red) and 123 proteins at lower level (yellow) in the MH proteome when compared to DME proteome, the abundance ratio >2.

**Supplementary Table S7:** 37 neuronal proteins were present at higher level in iERM and/or MH samples compared to DME and their 90 interactors found in our vitreous proteome analysis.

**Supplementary Table S8:** Significantly differed proteins between iERM and MH proteomes, q-value < 0.05. 53 proteins were present at higher level (red) and 65 proteins at lower level (yellow) in the iERM proteome when compared to MH proteome, the abundance ratio >2.

